# Adaptation of plasticity to predicted climates in Australian rainbowfishes (*Melanotaenia*) across climatically defined bioregions

**DOI:** 10.1101/859769

**Authors:** Jonathan Sandoval-Castillo, Katie Gates, Chris J. Brauer, Steve Smith, Louis Bernatchez, Luciano B. Beheregaray

## Abstract

Resilience to environmental stressors due to climate warming is influenced by local adaptations, including the capacity for plastic responses. The recent literature has focussed on genomic signatures of climatic adaptation, however little work has been done to address how plastic capacity may be influenced by biogeographic history and evolutionary processes. Here, we investigate phenotypic plasticity as a target of climatic selection, hypothesising that lineages that evolved under warmer climate will exhibit greater plastic adaptive resilience to thermal stress. This was tested using common garden experiments to compare gene expression regulation within and among a temperate, a subtropical and a desert ecotype of Australian rainbowfish. Individuals from each ecotype were subjected to contemporary and projected summer thermal conditions for 2070, and their global patterns of gene expression were characterized using liver transcriptomes. Critical thermal maximums were also determined for each ecotype to assess thermal tolerance. A comparative phylogenetic expression variance and evolution model framework was used to assess plastic and evolved changes in gene expression. Similar changes in both the direction and the magnitude of expressed genes were found within ecotypes. Although most expressed genes were identified in all ecotypes, 532 genes were identified as candidates subject to ecotype-specific directional selection. Twenty-three of those genes showed signal of adaptive (i.e. genetic-based) plastic response to future increases in temperature. Network analyses demonstrated centrality of these genes in thermal response pathways, along with several highly conserved hub genes thought to be integral for heat stress responses. The greatest adaptive resilience to warming was shown by the subtropical ecotype, followed by the desert and temperate ecotypes. Our findings indicate that vulnerability to climate change will be highly influenced by biogeographic factors, and we stress the need for integrative assessments of climatic adaptive traits for accurate estimations of population and ecosystem responses.

## Introduction

Characterizing mechanisms underpinning variation in ecological adaptation can assist in identifying biogeographic patterns of vulnerability and resilience to environmental change. Climate change has promoted numerous range shifts and local extinctions due to exposure of populations to conditions outside their zones of tolerance (Grabherr, Gottfried, & Pauli, 2009; Parmesan et al., 1999; Thomas & Lennon, 1999; Wiens, 2016). However, it is expected that some populations will be able to persist *in situ* if they are not already living at the edge of their tolerance limits; or, if they are able to acclimatise or adapt outside their current range of tolerance (Catullo, Ferrier, & Hoffmann, 2015; Hoffmann & Sgro, 2011; Stillman, 2003; Sunday, Bates, & Dulvy, 2011, 2012). Species’ distributions are strongly influenced by thermal conditions in their native climates; it is expected that tolerance ranges and vulnerability to change will also be influenced by biogeographic factors (Addo-Bediako, Chown, & Gaston, 2000; Calosi, Bilton, Spicer, Votier, & Atfield, 2010; Cohet, Vouidibio, & David, 1980; Compton, Rijkenberg, Drent, & Piersma, 2007). Exploring how molecular mechanisms influence thermal resilience and, ultimately, the evolution of divergent thermal phenotypes, is an important step for inferring responses to a warming environment (Komoroske, Connon, Jeffries, & Fangue, 2015). While evidence suggests that plastic regulation of gene expression plays an important role in ecological adaptation, the effects of selection on plasticity are so far poorly understood and successfully untangling them is likely to require integrative approaches (Gilad, Oshlack, & Rifkin, 2006; Jin et al., 2001; McCairns & Bernatchez, 2012).

Plasticity can be described as a change in expressed phenotype as a function of the environment, and occurs through direct effects of the environment on allelic expression, as well as changes in interactions among loci (Nonaka, Svanbäck, Thibert-Plante, Englund, & Brännström, 2015; Scheiner, 1993). Here, we focus on plasticity as the ability or tendency of an individual to up- or down-regulate genes in response to the environment. For many genes, this occurs primarily at the level of transcription, and a complexity of plastic responses (i.e. adaptive, maladaptive or neutral) have been observed in regard to individual fitness (Ghalambor, McKay, Carroll, & Reznick, 2007; Gibert, Debat, & Ghalambor, 2019; Marden, 2008). For instance, plasticity can act as a buffer against environmental pressures, providing short-term advantage but potentially dampening the effects of natural selection (Ghalambor et al., 2007; Grenier, Barre, & Litrico, 2016). Plastic responses have also been found to increase the potential for colonisation of new areas and provide greater likelihood of adaptive radiation (Muschick, Barluenga, Salzburger, & Meyer, 2011; Pfennig et al., 2010; Wellband & Heath, 2017; Wund, 2012). Plasticity itself can be a target of selection if genotypes differ in their sensitivity to environmental variation (Fusco & Minelli, 2010), including selection driving rapid adaptive evolution of genes exhibiting non-adaptive plasticity (Ghalambor et al., 2015).

In the context of climate, studies of gene expression can inform about the functional pathways relevant for persistence under a given condition, as well as the likely targets of selection (Komoroske et al., 2015; Reed, Schindler, & Waples, 2011). This is especially important where phenotypes of ecological relevance are not obvious, and may be difficult to distinguish using traditional approaches (Nevins & Potti, 2007; Nosil, 2012). While a variety of methods have been developed to detect evidence of ecological adaptation at the genomic level, relatively few studies have so far attempted to find signals of selection acting on gene expression. Challenges include controlling for the large range of internal and external environmental variables influencing expression (Conesa et al., 2016; De Wit et al., 2012), as well as the effects of genetic distance which are typically expected to account for much of the variation in transcription observed between lineages (Dunn, Luo, & Wu, 2013).

Climatically defined bioregions provide a scale at which environmental variation drives meaningful differences in evolutionary and ecological processes (Fine, 2015; Jetz & Fine, 2012). The ability of populations to persist under climate warming is predicted to vary geographically (Aitken & Whitlock, 2013; Catullo et al., 2015; Ghalambor, Huey, Martin, Tewksbury, & Wang, 2006; Polato et al., 2018; Sorte, Jones, & Miller, 2011; Thomas et al., 2004), making climatic bioregions valuable systems for comparative studies of adaptation. For instance, the climatic variability hypothesis (CVH) predicts a positive relationship between breadth of thermal tolerance and the level of climatic variability experienced by organisms as latitude increases (Janzen, 1967). Studies of climate change impacts are increasingly seeking to integrate spatial modelling (e.g. climatic envelopes) to uncover associations between landscape features and evolutionary processes such as temperature adaptation (Araújo, Whittaker, Ladle, & Erhard, 2005; Cianfrani, Satizábal, & Randin, 2015; Summers, Bryan, Crossman, & Meyer, 2012). While a majority of species distribution models are primarily correlative, there has been an urgent call for an increase in mechanistic approaches for predicting species’ responses to climate change (Bay et al., 2017; Cavaleri, Reed, Smith, & Wood, 2015; Comte & Olden, 2016; Evans, Merow, Record, McMahon, & Enquist, 2016; Mathewson et al., 2017). Mechanistic approaches have the advantage of explaining the underlying processes associated with observed trends, minimising the risk of flawed extrapolation (Dormann et al., 2012; Kearney & Porter, 2009). On a molecular scale, this is important for disentangling the respective effects of adaptation and phylogenetic signal, as well as the interactions between these two factors (Comte & Olden, 2016; Hoffmann, Chown, & Clusella-Trullas, 2013).

Freshwater ecosystems are of particular interest when studying the impacts of climate change due to their vulnerability. They are dependent on the interrelated influences of temperature and precipitation, and their spatially fragmented and often linear nature reduces opportunities for organismal dispersal. Freshwater fishes represent an important component of vertebrate diversity and, as ectotherms, are especially vulnerable to thermal changes (Deutsch et al., 2008). The subjects of this study are Australian rainbowfishes of *Melanotaenia* (family *Melanotaeniidae*), a freshwater genus with historical origins in tropical southern New Guinea (McGuigan, Zhu, Allen, & Moritz, 2000). *Melanotaenia* spp. of the ‘australis’ clade (Unmack *et al*. 2013) provide an ideal model system to study climatic-driven adaptive evolution and to address predictions from the CVH for freshwater ecosystems. The clade contains a minimum of eight largely allopatric species that recently radiated into tropical, subtropical, desert and temperate regions of mainland Australia (Unmack *et al*. 2013). Species of this clade show adaptive phenotypic divergence due to selection linked to the hydrological environment (McGuigan, Chenoweth, & Blows, 2005; McGuigan, Franklin, Moritz, & Blows, 2003), as well as adaptive genomic divergence associated with hydroclimatic variation (Brauer, Unmack, Smith, Bernatchez, & Beheregaray, 2018; McGuigan et al., 2005; McGuigan et al., 2003) In terms of gene expression, common garden experiments in a subtropical ‘australis’ species (*M. duboulayi*) have tested the effect of 2070-projected summer temperatures on short-term (Smith, Bernatchez, & Beheregaray, 2013) and long-term (McCairns, Smith, Sasaki, Bernatchez, & Beheregaray, 2016) transcriptional responses. Both studies indicated a capacity for plastic response to future climates, and enabled identification of candidate genes for thermal adaptation (McCairns et al., 2016; Smith et al., 2013). In addition, the transgenerational experiment in *M. duboulayi* revealed pedigree-based evidence for heritability of observed plastic responses (McCairns et al., 2016).

Our work focuses on three closely related ‘australis’ species, *M. splendida tatei, M. fluviatilis* and *M. duboulayi*. Their ranges show a striking concordance with three major contemporary climatic bioregions of the Australian continent (Fig. 1), suggesting that their evolution has been influenced by selective pressures associated with climatic regimes. For this reason, we refer to them herein as climatic ‘ecotypes’, *sensu* Engelhard, Ellis, Payne, ter Hofstede, and Pinnegar (2010). We used an experimental approach to compare short-term transcriptional responses to a projected future temperature in subtropical, temperate and desert rainbowfish ecotypes. In addition, physiological tolerance to thermal stress was assessed by empirically estimating the critical thermal maximum of each ecotype. We hypothesise that ecotype resilience in future climates will be dependent on the biogeographic region in which a given ecotype has evolved. As such, we also predict to find evidence for adaptation of plastic responses to temperature among ecotypes. To test this, we applied a comparative phylogenetic expression variance and evolution model framework to detect transcriptional responses subject to ecotype-specific directional selection. This enabled us to explore how divergent selection on gene expression may have contributed to differences in thermal tolerance and to adaptive evolution in these climatically defined ecotypes.

**Fig. 1.**
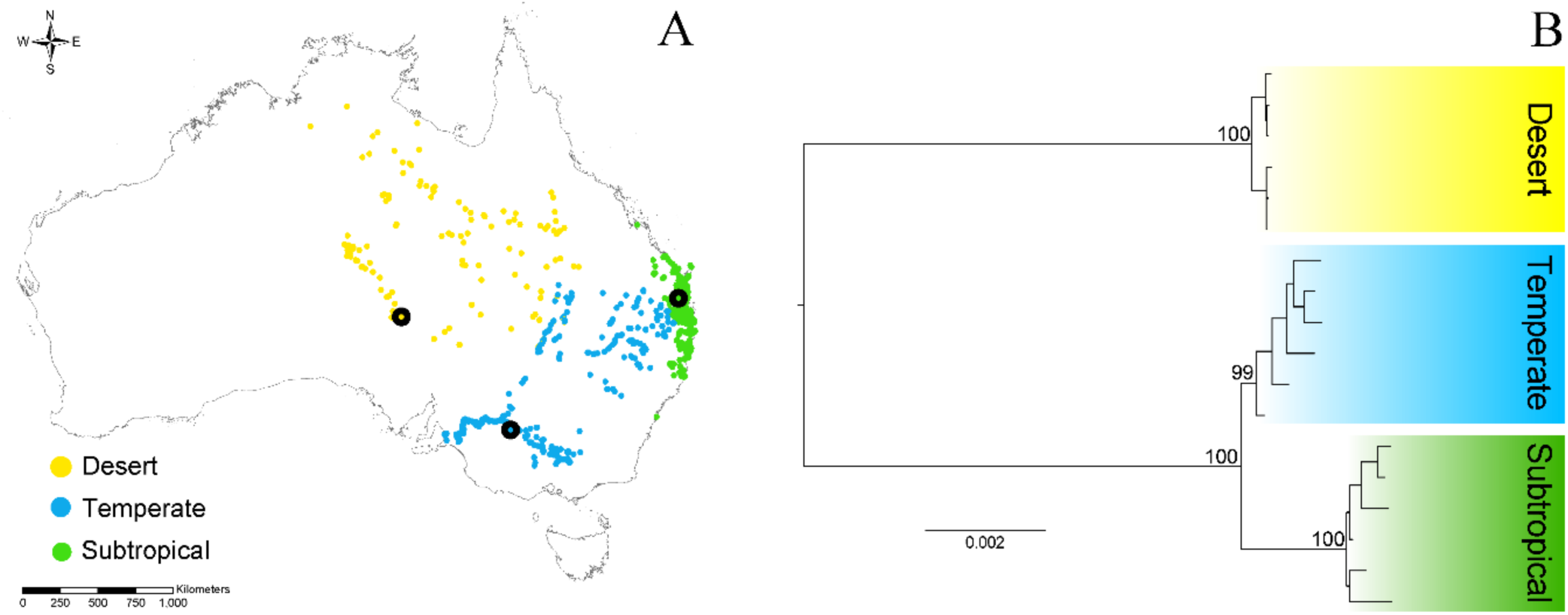
**A)** Spatially-validated records and range of the three *Melanotaenia* ecotypes, black circles show sampling localities for the transcriptomic and physiological experiments. **B)** Maximum Likelihood tree depicting evolutionary relationships among 36 individuals of the three ecotypes based on ddRAD sequences of 529 loci and 44681 bp. Numbers above nodes denote bootstrap support values.

## Methods

### Ecotype range, sampling and temperature experiments

The evolution of divergent expression of genes related to thermal tolerance was assessed in three closely related species of Australian rainbowfishes (Unmack *et al*. 2013). These are the Crimson spotted rainbowfish (*Melanotaenia duboulayi*) – a species with a subtropical distribution along coastal catchments of eastern Australia, the Murray River rainbowfish (*M. fluviatilis*) – a temperate species found in the inland Murray-Darling Basin, and the desert rainbowfish (*M. splendida tatei*) – a species found in arid and semi-arid catchments of central Australia (Figure 1). *Melanotaenia duboulayi* individuals were collected using bait traps and hand nets from the upper section of the Brisbane River, near the township of Fernvale in Queensland (subtropical; 27°26’37.39”S, 152°40’12.76”E). *Melanotaenia fluviatilis* individuals were collected from the mid section of the Murray River, close to the town of Gol Gol in New South Wales (temperate; 34°10’50.3”S 142°13’16.8”E) using a seine net. *Melanotaenia splendida tate* individuals were collected from Algebuckina Waterhole in South Australia (desert; 27°51’53.9”S 135°53’57.1”E) using fyke nets. Between 42 and 60 individuals were collected at each locality. The fish were transported live to the Flinders University animal rearing facility and acclimatised at 21°C for a minimum of 60 days prior to the start of temperature trials. Individuals from each species were maintained in single sex aquaria (∼20 fish/100L) at 21°C under conditions of 12 h light/12 h dark and fed once daily on a mixture of blood worms and fish pellets. To assess short-term responses to contemporary (21°C) and 2070-projected (33°C) average Australian summer temperatures, individuals of each species were randomly assigned to a treatment or a control group (n = 6 per group, per species). Temperature in these ‘climate-change treatment’ groups was increased by 2°C per day over a period of six days towards the target of 33°C, and then maintained for 14 days. The temperature (33°C) is the projected average summer temperature for Australia’s east coast in 2070 based on a high emission scenario (RCP8.5) of the International Panel on Climate Change (CSIRO, 2016; Smith et al., 2013). Control groups were kept at 21°C for the duration of the experiment. Fish were euthanized in an overdose of AQUI-S^®^ solution (50% isoeugenol) and immediately dissected to extract the liver. Sampling procedures took place in the same period of the day, between 9:00 am and 11:00 am. Only adult males of similar length were used to control for sex and age-related effects on transcription responses. Liver tissue was incubated at 4°C for 12 hr in RNAlater (Ambion) as per manufacturer’s instructions before storage at −80°C. In addition to being a relatively homogenous tissue, liver was selected because metabolic conditioning and gene expression is known to respond to heat stress (McCairns et al., 2016; Smith et al., 2013).

### RNA extraction, Illumina libraries preparation and sequencing

Total RNA was extracted from each individual liver tissue sample using the Ambion Magmax™-96 total RNA isolation kit (Life Sciences) according to the manufacturer’s instructions and following Smith et al (2013). Integrity and concentration were evaluated with an RNA Nano assay kit on an Agilent Bioanalyzer 2100 (Agilent Technologies) and purity assessed using a NanoDrop 1000 spectrophotometer (Thermo Scientific). Normalised quantities of total RNA were then used to prepare 36 separate Illumina sequencing libraries with the TruSeq™ RNA sample preparation kit (Illumina) using the adapter indices supplied by the manufacturer (Illumina MID tags 2, 4–7, 12–16, 18, 19) and following Gates, Sandoval-Castillo, Bernatchez, and Beheregaray (2017). Individual libraries were normalised and pooled together in groups of 12 samples. The resulting three pools were each sequenced in separate 100 bp paired-end lanes in an Illumina HiSeq2000 at Génome Québec Innovation Centre in Montreal, Canada.

### Read filtering, *de novo* assembly and annotation

Sequence data were demultiplexed by individual and trimmed of indexing adaptors. Low-quality (Q<20) bases were trimmed, then remnant adapter sequences, low quality reads, and reads shorter than 45 bp were removed using TRIMMOMATIC V0.36 (Bolger, Lohse, & Usadel, 2014). Four transcriptomes were assembled *de novo* with the program TRINITY V2.5.1 (Grabherr et al., 2011) using a pipeline described in Gates et al. (2017). One transcriptome for each ecotype was assembled using both experiment and control groups of each ecotype, then a *Melanotaenia* transcriptome was assembled using the samples of all ecotypes combined. The success of the completed transcriptome assembly was evaluated using read content statistics (% raw reads present), contig length distribution (N50), annotation-based metrics (% full length transcripts) and Benchmarking Universal Single-Copy Orthologs (BUSCO) software (Simão, Waterhouse, Ioannidis, Kriventseva, & Zdobnov, 2015) (Table S1). The open reading frames (ORFs) of a minimum length of 100 peptides were extracted from the assembled *Melanotaenia* transcriptome using the script TRANSDECODER V3.0 (Haas et al., 2013) and identified as candidate protein coding regions. Then, where two or more transcripts showed 80% or higher similarity, all but the longest transcript were removed to generate a non-redundant set of transcripts; herein referred to as ‘unigenes’. The whole transcriptome was functionally annotated using the command blastx (Altschul, Gish, Miller, Myers, & Lipman, 1990) to query uni-genes against the UniprotKB protein database (using a 1 x 10^−2^ *e*-value cut-off; Consortium, 2014) to identify homology to known proteins. In addition, any transcript showing 50% or higher similarity to any bacteria, fungus or virus genes were removed from further analysis.

### Transcript quantification and differential expression analysis

To test for differential expression (DE) between experimental groups and among ecotypes, reads for each sample were mapped back to the predicted protein coding regions using BOWTIE2 V2.2.7 (Langmead & Salzberg, 2012), then gene-level abundance estimations were performed with RSEM V1.2.19 (Li & Dewey, 2011). To enable comparison of expression level among samples, read count estimations were cross-sample-normalized using the trimmed mean of M-values method (TMM; Robinson and Oshlack, 2010). Normalized count data were then used as input for the program DESeq2 V1.10.1 (Love, Huber, & Anders, 2014). We used a conventional threshold (e.g. McCulloch et al., 2019) specifying that transcripts with a minimum log2 fold change of two between any two groups (i.e. experiment vs control, ecotype vs ecotype) were considered differentially expressed at a false discovery rate of 5%.

### Gene expression plasticity and divergent selection

We implemented the Expression Variance and Evolution Model (EVE) (Rohlfs & Nielsen, 2015) using the transcriptome of three ecotypes of *Melanotaenia* to identify transcripts potentially under divergent selection for expression levels. Briefly, the model uses a phylogenetic tree and expression data to estimate a parameter β that represents the ratio of among-lineage expression divergence to within-lineage expression diversity. This ratio should be approximately constant over most genes if no divergent selection is acting between lineages. For each transcript (i), the EVE model assesses the null hypothesis that independent transcript β_i_ is not significantly different to a shared β_s_ for all transcripts; if β_i_ is higher than β_s,_ the model assumes that transcript i is subject to lineage-specific directional selection on expression level. Following Brauer, Unmack, and Beheregaray (2017), we considered transcripts to be under divergent selection when βi was significantly higher than βs at a false discovery rate of 10%.

To calculate the expected expression covariance between lineages under shared and independent evolutionary history scenarios, we constructed a phylogenetic tree, using genome-wide SNP (single nucleotide polymorphisms) data from 12 samples of each ecotype. These data were obtained using reduced-representation sequencing methods (ddRAD) in previous studies of population genomics of the three ecotypes (Brauer et al. 2018, Attard et al. in review; Smith et al. in review; Supplementary Information X). The software PyRAD V3.0.6 (Eaton, 2014) was used to align the genome-wide sequences and RAXML V8.2.1 (Stamatakis, 2014) was used to perform a maximum-likelihood phylogenetic analysis, with the GTRGAMMA model and 1000 bootstrap replicates. The final concatenated dataset for the 36 rainbowfishes was based on 529 loci and 44681 bp. The consensus phylogenetic tree was used as the input phylogeny for the EVE analysis.

### Gene ontology enrichment analysis and pathway network analysis

We performed an enrichment analysis on the DE genes and on the EVE candidate genes relative to all genes, using the R package TOPGO v2.34 (Alexa & Rahnenfuhrer, 2010). For this analysis, terms were considered to be enriched if they were significant in both Fisher’s classic and weight tests with a P=<0.01. Moreover, to understand the relative importance of candidate and shared plastic genes in the response of rainbowfishes to heat stress, a network analysis was conducted using CYTOSCAPE V3.7 (Shannon et al., 2003). First, a protein interaction network was created from the entire DE gene set by drawing edges between genes with physical and functional interactions reported for humans and with orthologous functions in zebrafish in the STRING database (Szklarczyk et al., 2016). The relative importance of a protein is correlated with its connectivity in an interactive network. We calculated the node degrees as an estimator of protein connectivity. Then we identified highly connected genes (hubs) as those with a node degree greater than or equal to the sum of the mean plus twice the standard deviation of the node degree distribution (Rakshit, Rathi, & Roy, 2014).

### Determination of thermal tolerance (CT_MAX_)

We determined the thermal tolerance of each ecotype via short-term CT_MAX_ experiments following Becker and Genoway (1979). To control for sex and age-related effects, we collected 10 females of each ecotype from the same populations used for the transcriptomic experiments. After acclimation to 21°C for a minimum period of 60 days, each fish was placed individually in a 5 L glass beaker containing 3L of water at 21°C. Water temperature was increased at a rate of approximately 1°C every 3 minutes (rate of 0.33°C/min) using a digital water bath SWBD (Stuart^®^). We stopped the experiment and recorded the temperature when the fish showed both motor disorganization and loss of equilibrium for a period of one minute. Motor disorganization was considered when the swimming pattern was evidently changed with respect to their original condition; loss of equilibrium was determined when the fish turned upside down while swimming. Thermal limit for a given ecotype was obtained by averaging over 10 independent replicates. An ANOVA test was used to assess statistical differences in CT_MAX_ among ecotypes.

## Results

### Transcriptome sequencing and assembly

Illumina sequencing of the 36 individual libraries produced over 848 million paired-end reads (2 x 100 bp). After trimming and quality filtering, 741 million reads (87.4%) were retained (Table S1) for the *de novo* assemblies of each of the three ecotypes as well as the combined assembly for the genus *Melanotaenia*. The *Melanotaenia* assembly resulted in a total of 457,235 contigs (‘Trinity transcripts’), and 269,386 genes (‘Trinity genes’) from which 37,160 ORFs were detected and 34,815 uni-genes were identified (Table S1). Based on all transcript contigs, an N50 of 1702 and an average contig length of 904 were achieved. Assembly completeness assessment using BUSCO found a high percentage of genes in common with fish gene datasets (e.g. 89.2% for *Melanotaenia*, Fig. S1). All downstream analyses were based on the reference set of 34,815 *de novo* assembled uni-genes.

### Differential expression analysis

Of the 34,815 uni-genes, over 81% (28,483) were present in all three *Melanotaenia* ecotypes, with low percentages of uni-genes exclusive to each ecotype (Fig. 2A). Comparison of gene expression profiles among ecotypes and between climate-change treatments identified 2,409 differentially expressed (DE) uni-genes. Expression profiles of these genes showed a strong phylogenetic pattern with individuals showing the highest correlation of transcription responses within ecotypes, followed by high correlation between experiment and control groups within each ecotype (Fig. 2B, see also below). On the other hand, when gene expression was compared between experimental treatments, only 236 DE uni-genes were identified (Fig. 3A). Of these, 16 uni-genes were DE between treatments in two or more ecotypes (Fig. 3B), indicating shared plasticity for those genes. In contrast, unique plastic responses to projected summer temperatures were observed for the temperate ecotype in 27 uni-genes, the desert ecotype in 84 uni-genes and the subtropical ecotype in a much higher 109 uni-genes. This indicates a strong effect of phylogeny on plastic gene expression but may also represent the effects of divergent selection and adaptation to unique climatic ecoregions.

**Fig. 2.**
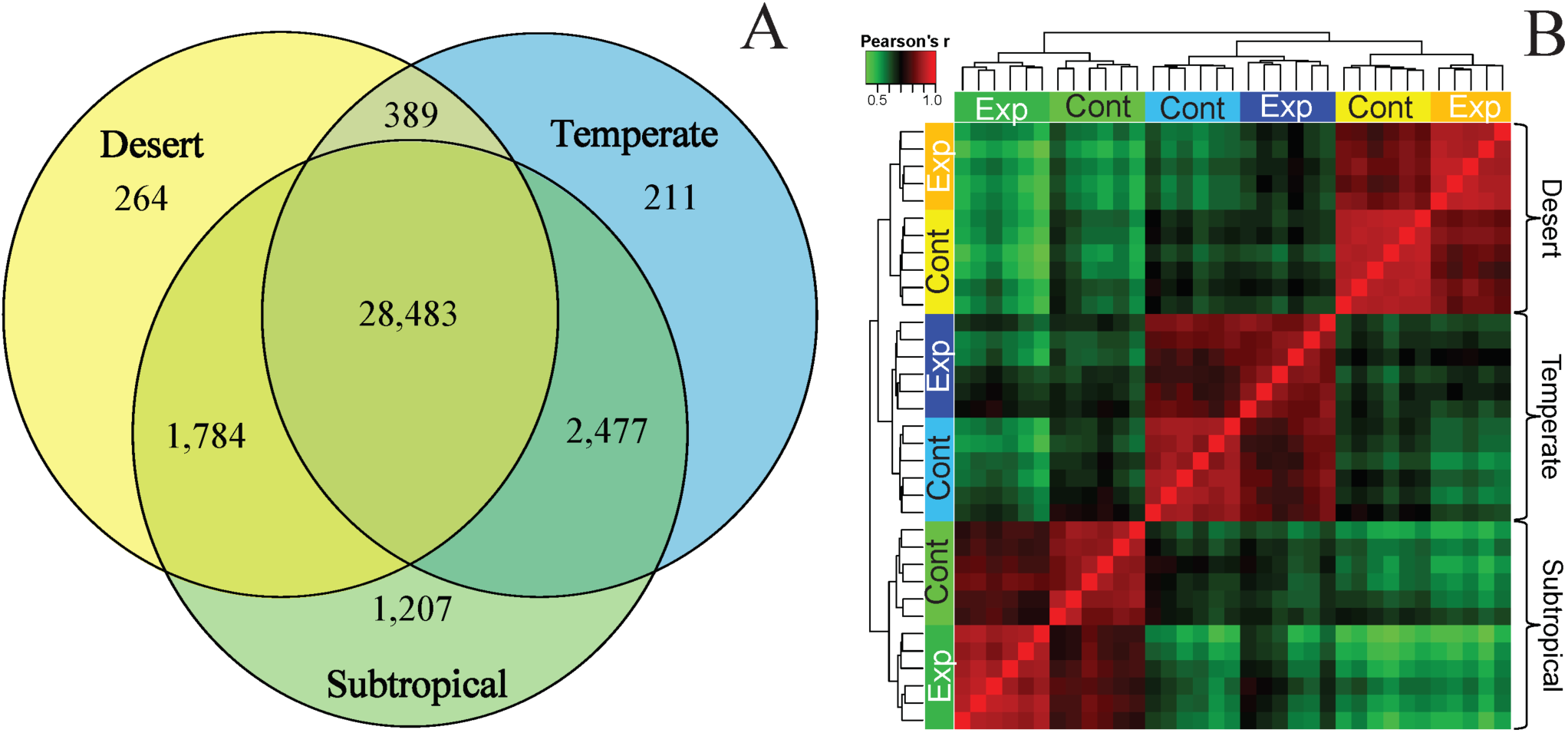
**A)** Venn diagram of uni-genes identified in each ecotype of *Melanotaenia* as well as shared among ecotypes (based on a total of 34,815 uni-genes). **B)** Heatmap summarizing correlation among ecotypes in log_2_ gene expression profiles. This analysis was based on 2,409 differentially expressed transcripts. Coloured bars under the sample dendrograms represent the climate-change experiment (Exp) and control (Cont) groups.

**Fig. 3.**
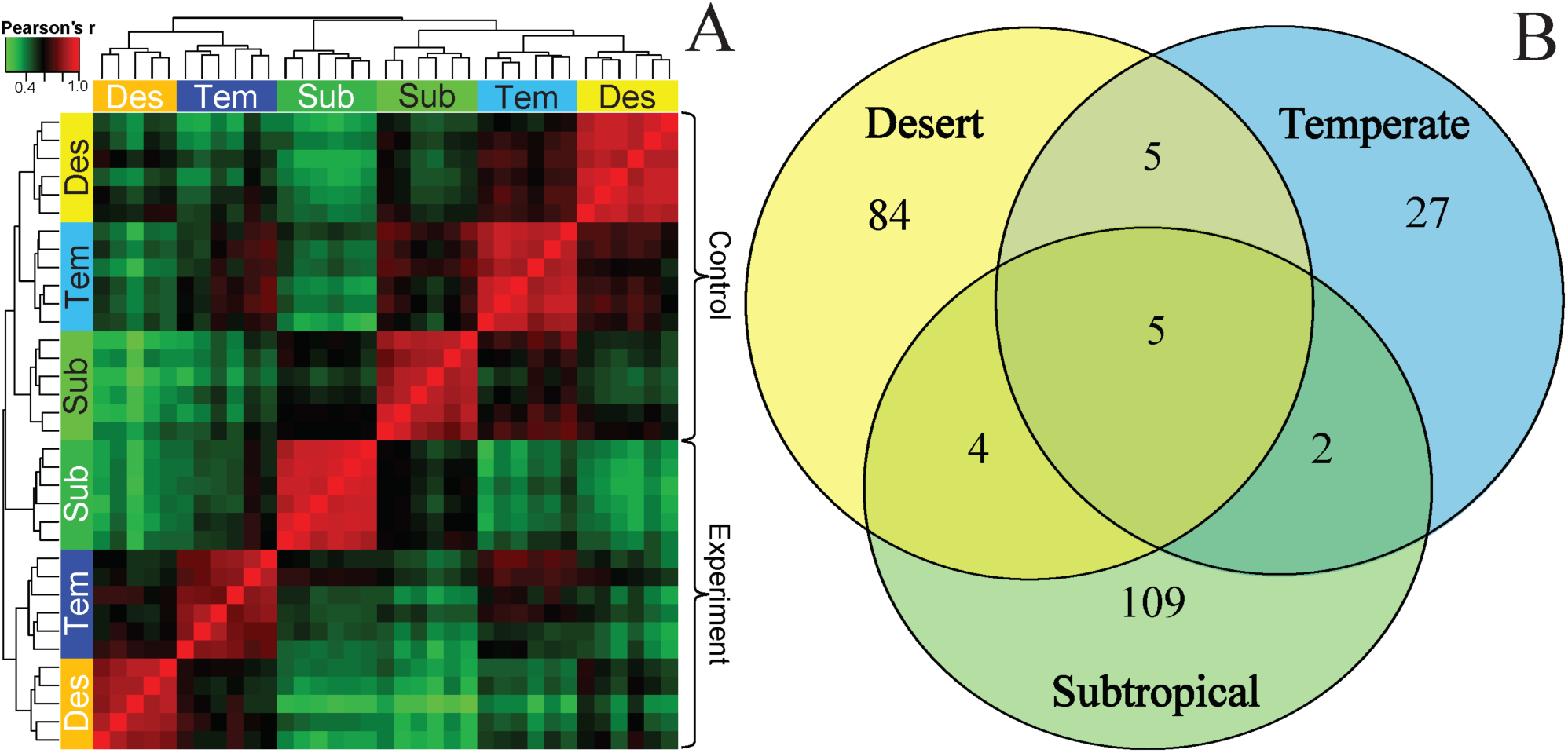
**A)** Heatmap summarizing correlation between treatments (Control vs Experiment) in log2 gene expression profiles. This analysis was based on 236 differentially expressed uni-genes identified between Control and Experiment samples. Coloured bars under the sample dendrograms represent the ecotypes, with climate-change experiment groups represented by dark colour variation and control groups represented by light colour variation. **B)** Venn diagram of differentially expressed uni-genes shared between ecotypes.

### Divergent selection on gene expression

The RAXML majority-rule consensus tree provided strong support for reciprocal monophyly of each ecotype (i.e. each named taxon), with all individuals from each ecotype clustering within a single clade (Fig. 1B). The topology is consistent with previous studies (McGuigan et al., 2000) that indicated a sister relationship between the temperate (*M. fluviatilis*) and the subtropical (*M. duboulayi*) ecotypes. This consensus tree was used as the input phylogeny for the EVE analysis (Fig. 1B). Of the 34,815 uni-genes assessed with the EVE, 532 were identified as candidates subject to lineage-specific directional selection on expression level (FDR 10%). These were the genes that showed greater expression variance among rather than within lineages after controlling for phylogenetic effects. The dendrogram of individuals based on these genes was consistent with phylogenetic patterns (Fig. S2) which showed greater differentiation between subtropical and desert ecotypes. Out of these 532 candidate genes, 23 were also identified as differentially expressed between treatments (Fig. 4A). The expression profiles of these genes are more similar between the same treatment for subtropical and temperate ecotypes, than between among treatments within the same ecotype. This was not the case for the desert ecotype, for which control and experiment are differentiated, yet cluster together. Only one of these 23 EVE candidate genes is DE between treatments in all ecotypes. This suggests that the plastic gene expression responses for these 23 genes are under divergent selection for resilience to heat stress among ecotypes, with the greatest differences between desert and the other two ecotypes.

**Fig. 4.**
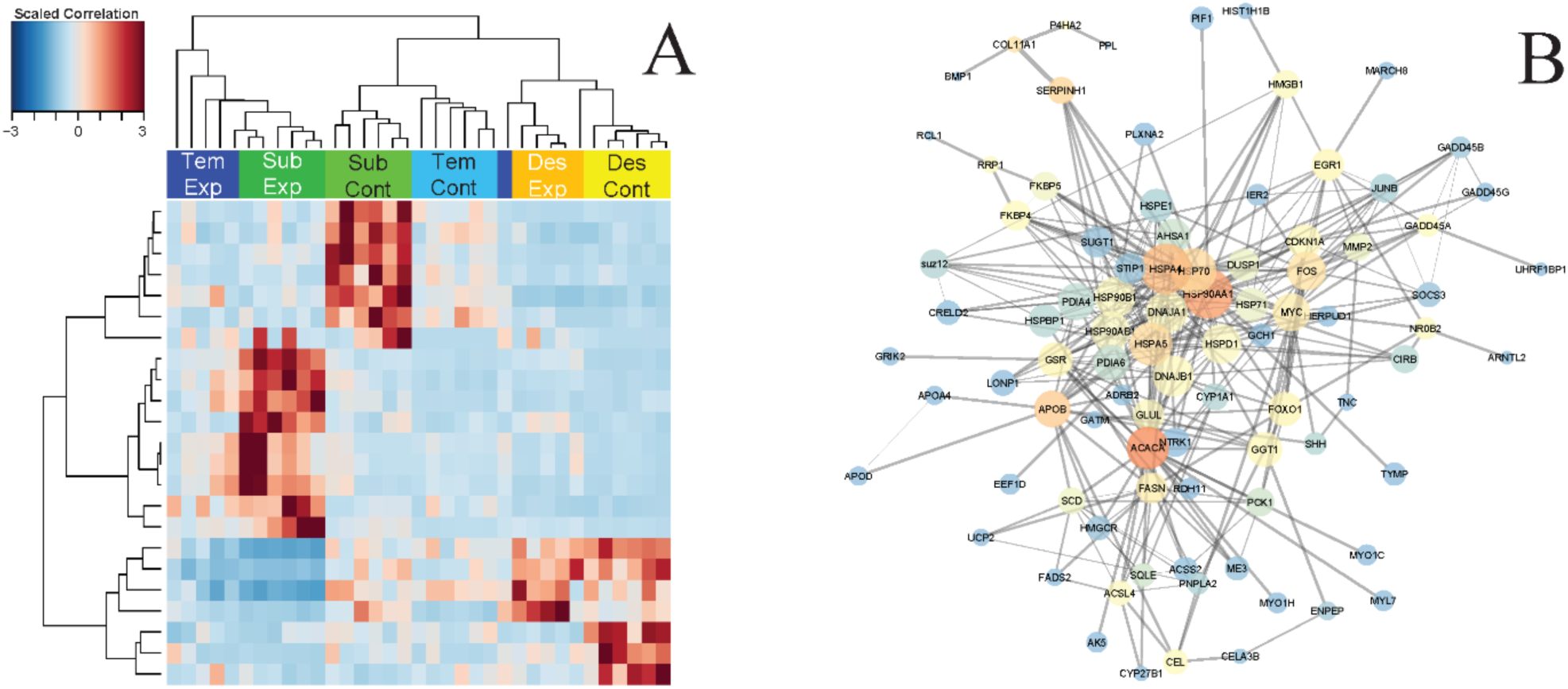
**(A)** Hierarchical clusters of 23 transcripts identified as candidates for divergent selection on expression level and also showing significant differential expression between control and experiment. Color bars indicate the ecotype of the samples. TemCont=Temperate control (20°C), TemExp= Temperate experimental (33°C), SubCont=Subtropical control (20°C), SubExp= Subtropical experimental (33°C), DesCont= Desert control (20°C), DesExp= Desert experimental (33°C). **(B)** Protein interaction network containing 137 heat stress associated proteins lined via 1114 interactions. Size of node is proportional to it centrality in the network, color of node indicate the relative number of interaction it is directly involved (blue lower – red higher number of interactions), both color and size of the node indicate realtive importance of the protein in *Melanotaenia* heat stress response.

### Functional annotation, enrichment analysis and network analysis

Uni-genes aligned to UniprotKB resulted in 25,315 protein hits, of which 24,276 (96%) were assigned to a total of 293,781 GO terms (Supplementary Information Table S6). Enrichment analysis of GO terms assigned to the 236 DE uni-genes between experiment and control (Fig. 3A) found terms for five molecular functions (MF), 13 biological processes (BP) and five cellular components (CC) significantly enriched (p < 0.01; Table S3). The same enrichment analysis using the 23 EVE candidates for divergent selection that were also identified as DE between treatments (Fig. 4A) found three MF, four BP and two CC terms significantly enriched (p<0.01; Table S4).

The protein network analysis identified six genes with high degree of interaction, all of which were heat shock proteins. These hub genes included the only uni-gene identified as DE in all three ecotypes, and as a candidate for divergent selection by the EVE model (Fig. 4B; Table S5). In addition, the 16 uni-gene candidates for shared plasticity found to be DE between treatments in two or more ecotypes, as well as the 23 EVE candidate genes related to heat stress response between treatments, showed higher average node degrees compared with the rest of the DE genes (Table S5). This suggests an important role of these genes in plastic and adaptive heat stress responses of rainbowfish, respectively.

### Empirical thermal tolerance (CT_MAX_)

Thermal tolerance was significantly different among ecotypes (p = <0.001, Table S2, Fig 5) with the highest CT_MAX_ shown by the subtropical ecotype (38.0°C; CI=37-38.6°C), followed by the desert ecotype (37.2°C; CI=36.1-37.6°C) and finally the temperate ecotype (34.9°C; CI=33.1-36.5°C). Interestingly, the inferred estimates of CT_MAX_ across ecotypes were correlated with the number of DE uni-genes between climate change treatments displayed by each ecotype (r= 0.998, Fig. 5).

**Fig. 5.**
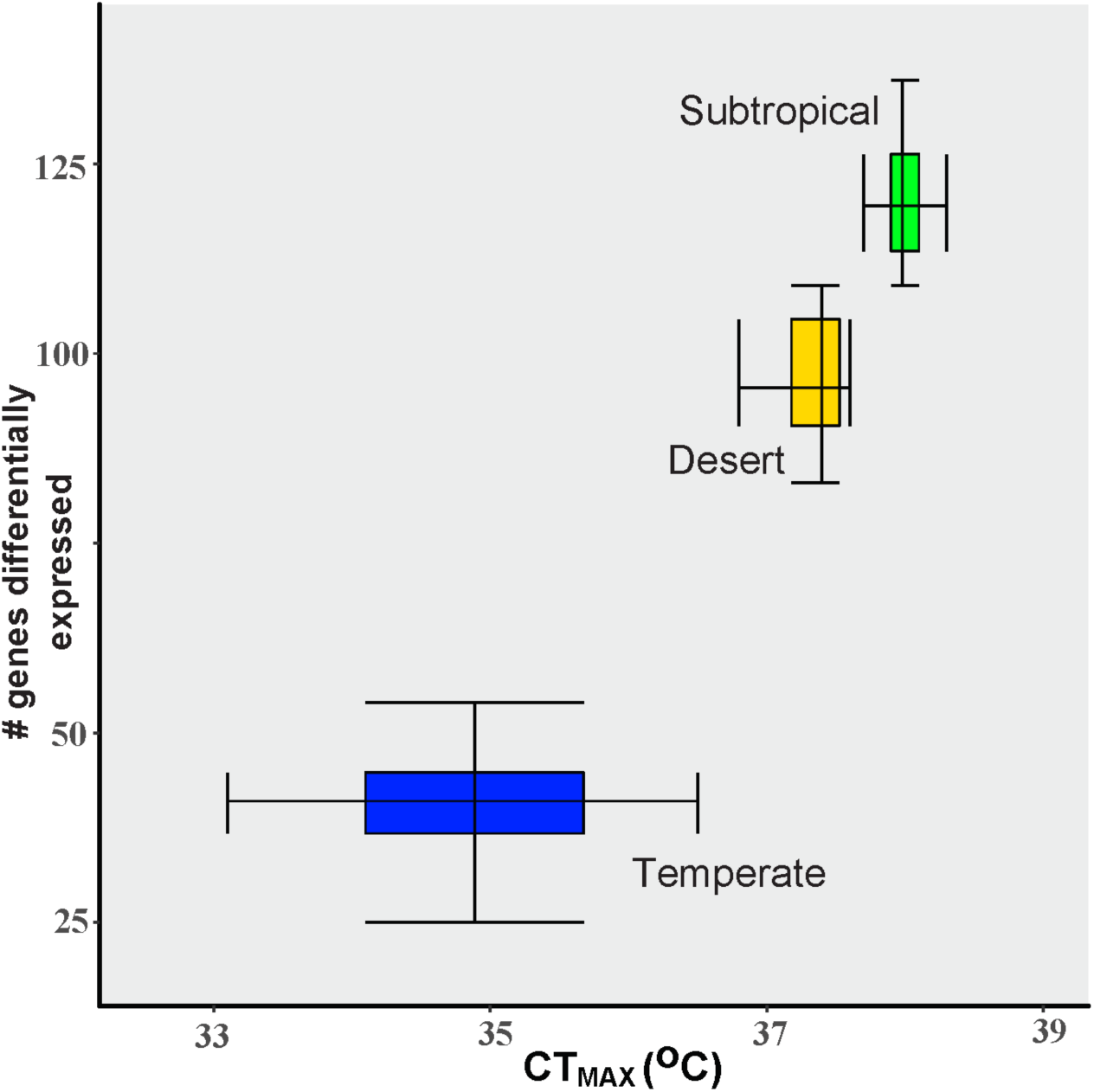
Association between CT_MAX_ and number of genes differentially expresed in response to heat stress in three ecotypes of *Melanotaenia* (r=0.998). Box plots display the uppper and lower quartiles, whiskers represent 95 and 5 percentiles, and their intersections represent the median.

## Discussion

We compared transcriptional plasticity to projected future temperatures and physiological tolerance to thermal stress in three contrasting climatic ecotypes of Australian rainbowfish: temperate, desert and subtropical. Within ecotypes, individuals exhibited very similar changes in both the direction and the magnitude of expressed genes. On the other hand, despite most expressed genes being identified in the transcriptomes of all ecotypes (i.e. 81% of the 34,815 uni-genes), response mechanisms to predicted thermal stress differed remarkably among ecotypes. Interestingly, both the plastic responses and tolerance to thermal increases varied in a biogeographically determined manner. Subtropical rainbowfish showed both the highest transcription response and tolerance to thermal stress amongst ecotypes, temperate rainbowfish showed the lowest responses for both mechanisms, and desert rainbowfish showed intermediate transcriptional responses and physiological tolerance. Although all species mounted substantial responses to 2070-projected summer temperatures, a striking result was that transcriptional changes common to all three ecotypes were limited to only five genes, thus confirming variation in plastic responses to thermal stress among ecotypes.

Such divergence in plastic responses between ecotypes is consistent with lineage-specific adaptation resulting from contrasting selective pressures across the climatically-defined bioregions, but can also be associated with neutral mechanisms of evolution (Dunn et al., 2013; Whitehead, 2012; Whitehead & Crawford, 2006). For this reason, we incorporated a phylogenetic model (EVE) to control for the effects of neutral drift on gene expression (Rohlfs & Nielsen, 2015). This approach identified a large suite of candidates for divergent selection on gene expression between ecotypes, of which a subset of 23 genes also showed significant ecotype-specific response to thermal manipulation (Fig. 4A). We consider these as strong candidates for adaptive (i.e. genetic-based) plastic response to future increases in temperature. Network analyses demonstrated centrality of these genes in thermal response pathways, while also identifying several highly conserved hub genes. These genes appear to be of fundamental importance for modulating thermal response pathways and adaptive potential in the three ecotypes. Together, these results show that while integral expression responses can be conserved among lineages, the tendency for divergence in response to thermal stress is high. This divergence not only exceeds neutral expectations but corresponds to differences in thermal tolerance across climatically defined bioregions, speaking to the importance of biogeographic history in considerations of climate-adaptive potential.

### Adaptive mechanisms contribute to gene expression plasticity among ecotypes

Shifts in gene expression regulation have for a long time been hypothesised to contribute to adaptive diversity (King & Wilson, 1975). However, the evolution of plastic responses by natural selection has been infrequently documented in empirical studies, particularly in natural populations (but see Whitehead and Crawford (2006); McCairns and Bernatchez (2010); Kenkel and Matz (2016); Roelofs et al. (2009); Kingsolver and Buckley (2017)). Although a diversity of mechanisms regulate gene expression (Orphanides & Reinberg, 2002), substantial empirical evidence supports heritability of expression responses (Roberge, Guderley, & Bernatchez, 2007; Schadt et al., 2003; Whitehead & Crawford, 2006), as recently demonstrated in the subtropical Australian rainbowfish (*M. duboulayi*) (McCairns et al., 2016). As such, plasticity is likely to be subject to the same broad evolutionary processes as other heritable traits. For instance, under directional selection, limited expression polymorphism is expected within ecotypes, while extensive divergence is expected between ecotypes (Hodgins-Davis & Townsend, 2009). Under stabilising selection, expression regulation is predicted to be highly consistent within and across ecotypes (Ghalambor et al., 2007; Hodgins-Davis & Townsend, 2009). Meanwhile, under neutral evolution, patterns of gene expression are expected to correlate with evolutionary divergence (Whitehead & Crawford, 2006), which in our case was assessed using a phylogenomic-based approach. Our comparative analyses suggested that all of the above mechanisms have influenced gene expression responses to projected thermal stress in rainbowfishes. This fits with our understanding of thermal tolerance adaptation in ectotherms as highly complex, and involving multiple levels of biological organisation (Cossins, 2012; Pritchard & Di Rienzo, 2010).

The majority of DE genes under 2070-projected temperature manipulation exhibited patterns of variation that could be associated with phylogenetic distance (Fig. 1B; Fig. 2B). This demonstrates that plastic responses to future climates can be highly constrained by demographic history, even among groups which are as recently diverged as Australian rainbowfishes (Unmack, Allen, & Johnson, 2013). Nonetheless, we were able to reject neutral scenarios as the most parsimonious explanation for the variation in a large subset of DE genes. In the case of the EVE candidates for directional selection to thermal stress, there was very little expression polymorphism within ecotypes, but high levels of divergence between ecotypes. We suggest that regulatory differences in these genes have helped to facilitate persistence of rainbowfish ecotypes within their respective thermal environments, bringing each closer to a local phenotypic optimum. While evidence of ecological selection on plasticity is rare, an example includes the gene expression divergence of a soil-dwelling hexapod (*Orchesella cincta*) in populations subsisting in contaminated mine sites (Roelofs et al., 2009). This was correlated with heavy metal tolerance, which resulted in a heritable increase in metal excretion efficiency. Similarly, Brennan, Galvez, and Whitehead (2015) demonstrated a shift in salinity-specific expression responses in populations of killifish (*Fundulus heteroclitus*) in populations adapted to habitats of contrasting salinity.

The centre of the gene interaction network for the three rainbowfish ecotypes consisted largely of heat shock proteins, which play an important role in thermal responses in a wide variety of taxa (Chen, Farrell, Matala, & Narum, 2018; Feder & Hofmann, 1999). Patterns of plastic responses to temperature were most likely to be shared among ecotypes in these central ‘hub’ genes (Table S5). This indicates a conserved functional role, which may have been retained through purifying and possibly stabilising selection. Hub genes influence the expression and activity of genes downstream in an expression network, and tend to be highly conserved in their expression between lineages (Evans, 2015). In genome-wide studies of model organisms, the deletion of hub genes is more likely to be deleterious than for non-hub genes. This can be due to either compromised network structure, or simply because they are more likely to be involved in essential interactions (He & Zhang, 2006). However, the fact that the EVE candidates for divergent expression among ecotypes also exhibited greater average connectedness than other DE genes (Table S5) suggests the importance of these genes in the respective ecological adaptations they have likely facilitated. In fact, three EVE candidates were also identified as hub genes, and one of these shows plasticity in all ecotypes (HSP90AA1). A change in expression in one or a few hub genes could therefore translate to a completely different stress response pathway. Indeed, enrichment analyses indicate that functions as diverse as metabolism, immune response, oxidative stress response, DNA damage response, signal transduction and other stress responses are contributing to local adaptation among temperate, desert and subtropical ecotypes.

While the transcriptomic methods used here directly address mechanisms for thermal response, we are not yet able to infer specific fitness effects of divergent expression patterns in warming climates. Despite this, the number of genes regulated in response to warming differed markedly between ecotypes, with the greatest number responding in the most heat-resilient subtropical ecotype (CT_MAX_ 38.0°), and the smallest number responding in the least heat-resilient temperate ecotype (CT_MAX_ 34.9°). Similarly, previous work comparing montane and desert redband trout (*Oncorhynchus mykiss gairdneri*) found that the more resilient desert trout regulated a larger number of genes than the less resilient montane trout in response to acute warming conditions (Garvin, Thorgaard, & Narum, 2015; Narum & Campbell, 2015). While the absolute number of transcripts regulated in a given condition can depend on many factors, including differences in constitutive expression or qualitative differences such as amino acid or regulatory changes (Somero, 2010), it is possible that a larger number of regulated genes may often reflect a more sophisticated plastic response to environmental stressors. In the case of rainbowfish, it is too early to say whether the observed increase in number of DE genes represents a more specialised adaptation to heat by the subtropical rainbowfish compared to desert and temperate ecotypes. However, the association between thermal tolerance and number of regulated transcripts does provide further evidence to support adaptive differences in the potential for expression-mediated phenotypic plasticity.

### Physiological thermal tolerance is specific to ecotype

It is generally assumed that organisms are adapted to or have the ability to acclimate to the temperatures normally encountered in their habitat range (Ghalambor et al., 2006). Thus, it has been proposed that organisms that evolved in warmer climates will have higher thermal tolerances than those in cool climates (Vernberg, 1959), and that those which have evolved in variable climates will have greater acclimation capacities and tolerance ranges than those in more stable climates (Janzen, 1967). Here, we found that critical thermal maximum (CT_MAX_) differed significantly among ecotypes, with the temperate ecotype displaying the lowest average thermal tolerance (34.9°C), the desert ecotype displaying intermediate thermal tolerance (37.2°C), and the subtropical ecotype displaying the highest thermal tolerance (38.0°C). Consistent with several studies assessing relationships between thermal tolerance and latitude (Addo-Bediako et al., 2000; Aguilar-Kirigin & Naya, 2013; Deutsch et al., 2008; Ghalambor et al., 2006; Magozzi & Calosi, 2015; Sunday et al., 2011), the observed pattern of rainbowfish CT_MAX_ is an increase with proximity to the equator. However, in this instance CT_MAX_ does not coincide with average maximum summer temperatures (or even average yearly temperatures) experienced in the climate of origin, with the hottest Australian temperatures found in the central deserts as opposed to the north-eastern subtropical region (Table S2, BOM 2014).

Perhaps counterintuitively, this finding is consistent with previous research which has emphasised the importance of temperature variability in relation to an organism’s upper limits of thermal tolerance (Ghalambor et al., 2006; Magozzi & Calosi, 2015; Sunday et al., 2011). Although wider ranges of tolerance have been found at higher latitudes, these have been largely attributed to lower critical thermal minimums of the temperate organisms studied (Addo-Bediako et al., 2000; Ghalambor et al., 2006). Meanwhile, higher thermal tolerances have been observed in tropical regions, but with an inverse relationship to tolerance breadth (Comte & Olden, 2016). This has led to the use of the term ‘climate specialists’ to describe tropical species, with an evolutionary trade-off suggested to exist between upper thermal tolerance and the capacity to acclimate to a wide range of temperatures (Comte & Olden, 2016; Deutsch et al., 2008; Payne & Smith, 2017; Tewksbury, Huey, & Deutsch, 2008).

Due to this apparent trade-off, our findings may highlight an unforeseen risk for desert taxa. The low humidity of desert environments allows rapid heat transfer, exposing organisms to some of the most extreme temperature variations both diurnally and annually. Of the three ecotypes assessed, the desert rainbowfish is currently faced with temperatures closest to its CT_MAX_ (Table S2; BOM, 2014), and its ability to adapt to extremely high temperatures may be compromised by the need to maintain a large window of tolerance. It is already common for ambient temperatures at the desert rainbowfish’s sampling location (BOM, 2014) to exceed its CT_MAX_, though larger water bodies are unlikely to reach such extremes due to the fluctuation in diurnal temperatures (∼15°C/day) and slow rates of heat exchange between air and water (Livingstone & Lotter, 1998). However, desert environments are predicted to experience more extreme heat waves and longer droughts under climate change scenarios (IPCC, 2014). This is likely to not only increase the length of time in which organisms are exposed to thermal stress conditions, but decrease the overall volume of aquatic refugia, making them more susceptible to ambient temperatures (Magoulick & Kobza, 2003). In such circumstances, typical behavioural responses such as seeking shade or cool-water sites created by deeper water or inflowing tributaries may be unable to compensate for these effects (Breau, Cunjak, & Peake, 2011; Magoulick & Kobza, 2003). Our data support a high risk for desert species relative to the other ecotypes, which is consistent with at least one species distribution modelling approach (Wiens, Stralberg, Jongsomjit, Howell, & Snyder, 2009) and a functional trait analysis (Vale & Brito, 2015), but not with a broader study using the vegetation sensitivity index (Seddon, Macias-Fauria, Long, Benz, & Willis, 2016). This lack of consensus is symptomatic of the current poor understanding of desert ecosystems and disparate approaches used to examine potential impacts of global change in these regions (Maestre, Salguero-Gomez, & Quero, 2012), highlighting the need for an integrated reassessment of dryland vulnerability to climate change.

### Conclusions and perspectives

Climate change is creating a discord between some organisms’ physiologies and their environments. In order to predict the likelihood of range shifts, population declines or local extinctions, it is useful to understand the distribution of adaptive diversity, including that of adaptive plasticity. However, despite extensive empirical studies about standing genetic variation and its effects on climate related traits, the concept of adaptive plasticity remains relatively unaddressed. Here, we compared transcriptional and physiological responses to projected future temperatures in three contrasting climatic ecotypes of Australian rainbowfish, to test the effects of ecotype-specific directional selection on plasticity. Our results supported the hypothesis that the capacity for plastic response to climate will vary biogeographically, even within a closely related group. Moreover, by controlling for the effects of phylogeny, we were able to infer that divergent selection on gene expression has contributed to observed differences in plastic capacity among ecotypes. By demonstrating immediate response mechanisms to thermal stress, as well as evidence for ecological selection on these mechanisms among lineages, our results emphasise the key role of plasticity in both short- and long-term climatic adaptation. While it is too early to link the observed transcriptional responses to direct measures of fitness, the proxy provided by the assessment of thermal tolerance provides a strong indication of ecological relevance. This has wide-ranging implications both for broad biogeographic assessments of climate impacts, as well as for more focussed predictions of species distribution changes which are only now beginning to account for intra-taxonomic adaptive variation. This study represents a stride towards a more holistic understanding of climatic adaptive potential in natural populations.

## Acknowledgements

We thank Leo O’Reilly, Catherine Attard and David Schmarr for assistance with sampling, and Leslie Morrison and her team for assistance with fish husbandry. Animal ethical approval was received from Flinders University (AWC E342 and AWC E429). This study was funded by the Australian Research Council (ARC grants DP110101207 and DP150102903 to L.B.B and L.B. and FT130101068 to L.B.B.).

## References

Addo-Bediako, A., Chown, S. L., & Gaston, K. J. (2000). Thermal tolerance, climatic variability and latitude. Proceedings of the Royal Society B: Biological Sciences, 267(1445), 739–745.

Aguilar-Kirigin, Á. J., & Naya, D. E. (2013). Latitudinal patterns in phenotypic plasticity: the case of seasonal flexibility in lizards’ fat body size. Oecologia, 173(3), 745–752. doi:10.1007/s00442-013-2682-z

Aitken, S. N., & Whitlock, M. C. (2013). Assisted gene flow to facilitate local adaptation to climate change. Annual Review of Ecology, Evolution, and Systematics, 44, 367–388.

Alexa, A., & Rahnenfuhrer, J. (2010). topGO: enrichment analysis for gene ontology. R package version, 2(0).

Altschul, S., Gish, W., Miller, W., Myers, E., & Lipman, D. (1990). 3.1 Sequence searches-challenges. Journal of Molecular Biology, 215, 403–410.

Araújo, M. B., Whittaker, R. J., Ladle, R. J., & Erhard, M. (2005). Reducing uncertainty in projections of extinction risk from climate change. Global Ecology and Biogeography, 14(6), 529–538.

Bay, R. A., Rose, N., Barrett, R., Bernatchez, L., Ghalambor, C. K., Lasky, J. R., … Ralph, P. J. T. A. N. (2017). Predicting responses to contemporary environmental change using evolutionary response architectures. The American Naturalist, 189(5), 463–473.

Becker, C. D., & Genoway, R. G. (1979). Evaluation of the critical thermal maximum for determining thermal tolerance of freshwater fish. Environmental Biology of Fishes, 4(3), 245.

Bolger, A. M., Lohse, M., & Usadel, B. (2014). Trimmomatic: a flexible trimmer for Illumina sequence data. Bioinformatics, btu170.

BOM. (2014). Bureau of Meteorology climate data online.

Brauer, C. J., Unmack, P. J., & Beheregaray, L. B. (2017). Comparative ecological transcriptomics and the contribution of gene expression to the evolutionary potential of a threatened fish. Molecular Ecology, 26(24), 6841–6856.

Brauer, C. J., Unmack, P. J., Smith, S., Bernatchez, L., & Beheregaray, L. B. (2018). On the roles of landscape heterogeneity and environmental variation in determining population genomic structure in a dendritic system. Molecular Ecology, 27(17), 3484–3497.

Breau, C., Cunjak, R. A., & Peake, S. J. (2011). Behaviour during elevated water temperatures: can physiology explain movement of juvenile Atlantic salmon to cool water? Journal of Animal Ecology, 80(4), 844–853.

Brennan, R. S., Galvez, F., & Whitehead, A. (2015). Reciprocal osmotic challenges reveal mechanisms of divergence in phenotypic plasticity in the killifish *Fundulus heteroclitus*. The Journal of Experimental Biology, 218(8), 1212–1222. doi:10.1242/jeb.110445

Calosi, P., Bilton, D. T., Spicer, J. I., Votier, S. C., & Atfield, A. (2010). What determines a species’ geographical range? Thermal biology and latitudinal range size relationships in European diving beetles (Coleoptera: Dytiscidae). Journal of Animal Ecology, 79(1), 194–204.

Catullo, R. A., Ferrier, S., & Hoffmann, A. A. (2015). Extending spatial modelling of climate change responses beyond the realized niche: estimating, and accommodating, physiological limits and adaptive evolution. Global Ecology and Biogeography, 24(10), 1192–1202.

Cavaleri, M. A., Reed, S. C., Smith, W. K., & Wood, T. E. (2015). Urgent need for warming experiments in tropical forests. Global Change Biology, 21(6), 2111–2121.

Chen, Z., Farrell, A. P., Matala, A., & Narum, S. R. (2018). Mechanisms of thermal adaptation and evolutionary potential of conspecific populations to changing environments. Molecular Ecology, 27(3), 659–674. doi:doi:10.1111/mec.14475

Cianfrani, C., Satizábal, H. F., & Randin, C. (2015). A spatial modelling framework for assessing climate change impacts on freshwater ecosystems: Response of brown trout (Salmo trutta L.) biomass to warming water temperature. Ecological Modelling, 313, 1–12. doi:http://dx.doi.org/10.1016/j.ecolmodel.2015.06.023

Cohet, Y., Vouidibio, J., & David, J. (1980). Thermal tolerance and geographic distribution: a comparison of cosmopolitan and tropical endemic Drosophila species. Journal of Thermal Biology, 5(2), 69–74.

Compton, T. J., Rijkenberg, M. J., Drent, J., & Piersma, T. (2007). Thermal tolerance ranges and climate variability: a comparison between bivalves from differing climates. Journal of Experimental Marine Biology and Ecology, 352(1), 200–211.

Comte, L., & Olden, J. D. (2016). Evolutionary and environmental determinants of freshwater fish thermal tolerance and plasticity. Global Change Biology, 23(2), 728–736.

Conesa, A., Madrigal, P., Tarazona, S., Gomez-Cabrero, D., Cervera, A., McPherson, A., … Mortazavi, A. (2016). A survey of best practices for RNA-seq data analysis. Genome Biology, 17, 13. doi:10.1186/s13059-016-0881-8

Cossins, A. (2012). Temperature biology of animals: Springer Science & Business Media.

CSIRO. (2016). Climate change in Australia: Australian climate futures tool. Retrieved from https://www.climatechangeinaustralia.gov.au/en/climate-projections/climate-futures-tool/introduction-climate-futures/

De Wit, P., Pespeni, M. H., Ladner, J. T., Barshis, D. J., Seneca, F., Jaris, H., … Palumbi, S. R. (2012). The simple fool’s guide to population genomics via RNA-Seq: an introduction to high-throughput sequencing data analysis. Molecular Ecology Resources, 12(6), 1058–1067. doi:10.1111/1755-0998.12003

Deutsch, C. A., Tewksbury, J. J., Huey, R. B., Sheldon, K. S., Ghalambor, C. K., Haak, D. C., & Martin, P. R. (2008). Impacts of climate warming on terrestrial ectotherms across latitude. Proceedings of the National Academy of Sciences, 105(18), 6668–6672.

Dormann, C. F., Schymanski, S. J., Cabral, J., Chuine, I., Graham, C., Hartig, F., … Schröder, B. (2012). Correlation and process in species distribution models: bridging a dichotomy. Journal of Biogeography, 39(12), 2119–2131.

Dunn, C. W., Luo, X., & Wu, Z. (2013). Phylogenetic Analysis of Gene Expression. Integrative and Comparative Biology, 53(5), 847–856. doi:10.1093/icb/ict068

Eaton, D. A. (2014). PyRAD: assembly of de novo RADseq loci for phylogenetic analyses. Bioinformatics, btu121.

Engelhard, G. H., Ellis, J. R., Payne, M. R., ter Hofstede, R., & Pinnegar, J. K. (2010). Ecotypes as a concept for exploring responses to climate change in fish assemblages. ICES Journal of Marine Science. doi:10.1093/icesjms/fsq183

Evans, M. E., Merow, C., Record, S., McMahon, S. M., & Enquist, B. J. (2016). Towards process-based range modeling of many species. Trends in Ecology & Evolution, 31(11), 860–871.

Evans, T. (2015). Considerations for the use of transcriptomics in identifying the ‘genes that matter’for environmental adaptation. Journal of Experimental Biology, 218(12), 1925–1935.

Feder, M. E., & Hofmann, G. E. (1999). Heat-shock proteins, molecular chaperones, and the stress response: evolutionary and ecological physiology. Annual Review of Physiology, 61(1), 243–282.

Fine, P. V. (2015). Ecological and evolutionary drivers of geographic variation in species diversity. Annual Review of Ecology, Evolution, and Systematics, 46, 369–392.

Fusco, G., & Minelli, A. (2010). Phenotypic plasticity in development and evolution: facts and concepts. Introduction. Philosophical Transactions of the Royal Society of London. Series B, Biological sciences, 365(1540), 547–556. doi:10.1098/rstb.2009.0267

Garvin, M. R., Thorgaard, G. H., & Narum, S. R. (2015). Differential expression of genes that control respiration contribute to thermal adaptation in redband trout (*Oncorhynchus mykiss gairdneri*). Genome Biology and Evolution, 7(6), 1404–1414.

Gates, K., Sandoval-Castillo, J., Bernatchez, L., & Beheregaray, L. B. (2017). De novo transcriptome assembly and annotation for the desert rainbowfish (*Melanotaenia splendida tatei*) with comparison with candidate genes for future climates. Marine Genomics, 35, 63–68.

Ghalambor, C. K., Hoke, K. L., Ruell, E. W., Fischer, E. K., Reznick, D. N., & Hughes, K. A. (2015). Non-adaptive plasticity potentiates rapid adaptive evolution of gene expression in nature. Nature, 525(7569), 372.

Ghalambor, C. K., Huey, R. B., Martin, P. R., Tewksbury, J. J., & Wang, G. (2006). Are mountain passes higher in the tropics? Janzen’s hypothesis revisited. Integrative and Comparative Biology, 46(1), 5–17. doi:10.1093/icb/icj003

Ghalambor, C. K., McKay, J. K., Carroll, S. P., & Reznick, D. N. (2007). Adaptive versus non-adaptive phenotypic plasticity and the potential for contemporary adaptation in new environments. Functional Ecology, 21(3), 394–407.

Gibert, P., Debat, V., & Ghalambor, C. K. (2019). Phenotypic plasticity, global change, and the speed of adaptive evolution. Current Opinion in Insect Science.

Gilad, Y., Oshlack, A., & Rifkin, S. A. (2006). Natural selection on gene expression. Trends in Genetics, 22(8), 456–461.

Grabherr, G., Gottfried, M., & Pauli, H. (2009). Climate effects on mountain plants. Nature, 369(6480), 448; 448.

Grabherr, M. G., Haas, B. J., Yassour, M., Levin, J. Z., Thompson, D. A., Amit, I., … Zeng, Q. (2011). Trinity: reconstructing a full-length transcriptome without a genome from RNA-Seq data. Nature Biotechnology, 29(7), 644.

Grenier, S., Barre, P., & Litrico, I. (2016). Phenotypic Plasticity and Selection: Nonexclusive Mechanisms of Adaptation. Scientifica, 2016, 7021701–7021701. doi:10.1155/2016/7021701

Haas, B. J., Papanicolaou, A., Yassour, M., Grabherr, M., Blood, P. D., Bowden, J., … Lieber, M. (2013). De novo transcript sequence reconstruction from RNA-seq using the Trinity platform for reference generation and analysis. Nature Protocols, 8(8), 1494.

He, X., & Zhang, J. (2006). Why do hubs tend to be essential in protein networks? PLoS Genetics, 2(6), e88.

Hodgins-Davis, A., & Townsend, J. P. (2009). Evolving gene expression: from G to E to G× E. Trends in Ecology & Evolution, 24(12), 649–658.

Hoffmann, A. A., Chown, S. L., & Clusella-Trullas, S. (2013). Upper thermal limits in terrestrial ectotherms: how constrained are they? Functional Ecology, 27(4), 934–949.

Hoffmann, A. A., & Sgro, C. M. (2011). Climate change and evolutionary adaptation. Nature, 470(7335), 479–485.

IPCC. (2014). Climate Change 2014–Impacts, Adaptation and Vulnerability: Regional Aspects: Cambridge University Press.

Janzen, D. H. (1967). Why Mountain Passes are Higher in the Tropics. The American Naturalist, 101(919), 233–249. doi:10.1086/282487

Jetz, W., & Fine, P. V. (2012). Global gradients in vertebrate diversity predicted by historical area-productivity dynamics and contemporary environment. PLoS Biology, 10(3), e1001292.

Jin, W., Riley, R. M., Wolfinger, R. D., White, K. P., Passador-Gurgel, G., & Gibson, G. (2001). The contributions of sex, genotype and age to transcriptional variance in Drosophila melanogaster. Nature Genetics, 29(4), 389.

Kearney, M., & Porter, W. (2009). Mechanistic niche modelling: combining physiological and spatial data to predict species’ ranges. Ecology Letters, 12(4), 334–350.

Kenkel, C. D., & Matz, M. V. (2016). Gene expression plasticity as a mechanism of coral adaptation to a variable environment. Nature Ecology and Evolution, 1, 0014. doi:10.1038/s41559-016-0014

King, M.-C., & Wilson, A. C. (1975). Evolution at two levels in humans and chimpanzees. Science, 188(4184), 107–116.

Kingsolver, J. G., & Buckley, L. B. (2017). Evolution of plasticity and adaptive responses to climate change along climate gradients. Proceedings of the Royal Society B: Biological Sciences, 284(1860), 20170386.

Komoroske, L. M., Connon, R. E., Jeffries, K. M., & Fangue, N. A. (2015). Linking transcriptional responses to organismal tolerance reveals mechanisms of thermal sensitivity in a mesothermal endangered fish. Molecular Ecology, 24(19), 4960–4981. doi:10.1111/mec.13373

Langmead, B., & Salzberg, S. L. (2012). Fast gapped-read alignment with Bowtie 2. Nature Methods, 9(4), 357.

Li, B., & Dewey, C. N. (2011). RSEM: accurate transcript quantification from RNA-Seq data with or without a reference genome. BMC Bioinformatics, 12(1), 323.

Livingstone, D. M., & Lotter, A. F. (1998). The relationship between air and water temperatures in lakes of the Swiss Plateau: a case study with pal\sgmaelig; olimnological implications. Journal of Paleolimnology, 19(2), 181–198.

Love, M. I., Huber, W., & Anders, S. (2014). Moderated estimation of fold change and dispersion for RNA-seq data with DESeq2. Genome Biology, 15(12), 550.

Maestre, F. T., Salguero-Gomez, R., & Quero, J. L. (2012). It is getting hotter in here: determining and projecting the impacts of global environmental change on drylands. Philosophical Transactions of the Royal Society B: Biological Sciences, 367(1606), 3062–3075.

Magoulick, D. D., & Kobza, R. M. (2003). The role of refugia for fishes during drought: a review and synthesis. Freshwater Biology, 48(7), 1186–1198.

Magozzi, S., & Calosi, P. (2015). Integrating metabolic performance, thermal tolerance, and plasticity enables for more accurate predictions on species vulnerability to acute and chronic effects of global warming. Global Change Biology, 21(1), 181–194.

Marden, J. (2008). Quantitative and evolutionary biology of alternative splicing: how changing the mix of alternative transcripts affects phenotypic plasticity and reaction norms. Heredity, 100(2), 111–120.

Mathewson, P. D., Moyer-Horner, L., Beever, E. A., Briscoe, N. J., Kearney, M., Yahn, J. M., & Porter, W. P. (2017). Mechanistic variables can enhance predictive models of endotherm distributions: the American pika under current, past, and future climates. Global Change Biology, 23(3), 1048–1064.

McCairns, R. J. S., Smith, S., Sasaki, M., Bernatchez, L., & Beheregaray, L. B. (2016). The adaptive potential of subtropical rainbowfish in the face of climate change: heritability and heritable plasticity for the expression of candidate genes. Evolutionary Applications, 9, 531–545. doi:10.1111/eva.12363

McCairns, R. S., & Bernatchez, L. (2010). Adaptive divergence between freshwater and marine sticklebacks: insights into the role of phenotypic plasticity from an integrated analysis of candidate gene expression. Evolution: International Journal of Organic Evolution, 64(4), 1029–1047.

McCairns, R. S., & Bernatchez, L. (2012). Plasticity and heritability of morphological variation within and between parapatric stickleback demes. Journal of Evolutionary Biology, 25(6), 1097–1112.

McCulloch, G. A., Oliphant, A., Dearden, P. K., Veale, A. J., Ellen, C. W., & Waters, J. M. (2019). Comparative transcriptomic analysis of a wing-dimorphic stonefly reveals candidate wing loss genes. EvoDevo, 10(1), 21. doi:10.1186/s13227-019-0135-4

McGuigan, K., Chenoweth, S. F., & Blows, M. W. (2005). Phenotypic divergence along lines of genetic variance. The American Naturalist, 165(1), 32–43.

McGuigan, K., Franklin, C. E., Moritz, C., & Blows, M. W. (2003). Adaptation of rainbow fish to lake and stream habitats. Evolution, 57(1), 104–118.

McGuigan, K., Zhu, D., Allen, G. R., & Moritz, C. (2000). Phylogenetic relationships and historical biogeography of melanotaeniid fishes in Australia and New Guinea. Marine and Freshwater Research, 51(7), 713–723. doi:http://dx.doi.org/10.1071/MF99159

Muschick, M., Barluenga, M., Salzburger, W., & Meyer, A. (2011). Adaptive phenotypic plasticity in the Midas cichlid fish pharyngeal jaw and its relevance in adaptive radiation. BMC Evolutionary Biology, 11, 116–116. doi:10.1186/1471-2148-11-116

Narum, S. R., & Campbell, N. R. (2015). Transcriptomic response to heat stress among ecologically divergent populations of redband trout. BMC Genomics, 16(1), 103.

Nevins, J. R., & Potti, A. (2007). Mining gene expression profiles: expression signatures as cancer phenotypes. Nature Reviews Genetics, 8(8), 601.

Nonaka, E., Svanbäck, R., Thibert-Plante, X., Englund, G., & Brännström, Å. (2015). Mechanisms by which phenotypic plasticity affects adaptive divergence and ecological speciation. The American Naturalist, 186(5), E126–E143. doi:doi:10.1086/683231

Nosil, P. (2012). Ecological speciation: Oxford University Press.

Orphanides, G., & Reinberg, D. (2002). A unified theory of gene expression. Cell, 108(4), 439–451.

Parmesan, C., Ryrholm, N., Stefanescu, C., Hill, J. K., Thomas, C. D., Descimon, H., … Tammaru, T. (1999). Poleward shifts in geographical ranges of butterfly species associated with regional warming. Nature, 399(6736), 579–583.

Payne, N. L., & Smith, J. A. (2017). An alternative explanation for global trends in thermal tolerance. Ecology Letters, 20(1), 70–77.

Pfennig, D. W., Wund, M. A., Snell-Rood, E. C., Cruickshank, T., Schlichting, C. D., & Moczek, A. P. (2010). Phenotypic plasticity’s impacts on diversification and speciation. Trends in Ecology & Evolution, 25(8), 459–467.

Polato, N. R., Gill, B. A., Shah, A. A., Gray, M. M., Casner, K. L., Barthelet, A., … Encalada, A. C. (2018). Narrow thermal tolerance and low dispersal drive higher speciation in tropical mountains. Proceedings of the National Academy of Sciences, 115(49), 12471–12476.

Pritchard, J. K., & Di Rienzo, A. (2010). Adaptation–not by sweeps alone. Nature Reviews Genetics, 11(10), 665.

Rakshit, H., Rathi, N., & Roy, D. (2014). Construction and analysis of the protein-protein interaction networks based on gene expression profiles of Parkinson’s disease. PloS One, 9(8), e103047.

Reed, T. E., Schindler, D. E., & Waples, R. S. (2011). Interacting effects of phenotypic plasticity and evolution on population persistence in a changing climate. Conservation Biology, 25(1), 56–63.

Roberge, C., Guderley, H., & Bernatchez, L. (2007). Genome-wide Identification of Genes under Directional Selection: Gene Transcription Qst Scan in Diverging Atlantic Salmon Subpopulations. Genetics, 177(2), 1011–1022.

Robinson, M. D., & Oshlack, A. (2010). A scaling normalization method for differential expression analysis of RNA-seq data. Genome Biology, 11(3), R25.

Roelofs, D., Janssens, T. K., Timmermans, M. J., Nota, B., Marien, J., Bochdanovits, Z., … Van Straalen, N. M. (2009). Adaptive differences in gene expression associated with heavy metal tolerance in the soil arthropod *Orchesella cincta*. Molecular Ecology, 18(15), 3227–3239.

Rohlfs, R. V., & Nielsen, R. (2015). Phylogenetic ANOVA: the expression variance and evolution model for quantitative trait evolution. Systematic Biology, 64(5), 695–708.

Schadt, E. E., Monks, S. A., Drake, T. A., Lusis, A. J., Che, N., Colinayo, V., … Cavet, G. (2003). Genetics of gene expression surveyed in maize, mouse and man. Nature, 422(6929), 297.

Scheiner, S. M. (1993). Genetics and evolution of phenotypic plasticity. Annual Review of Ecology and Systematics, 24(1), 35–68.

Seddon, A. W., Macias-Fauria, M., Long, P. R., Benz, D., & Willis, K. J. (2016). Sensitivity of global terrestrial ecosystems to climate variability. Nature, 531(7593), 229.

Shannon, P., Markiel, A., Ozier, O., Baliga, N. S., Wang, J. T., Ramage, D., … Ideker, T. (2003). Cytoscape: a software environment for integrated models of biomolecular interaction networks. Genome Research, 13(11), 2498–2504.

Simão, F. A., Waterhouse, R. M., Ioannidis, P., Kriventseva, E. V., & Zdobnov, E. M. (2015). BUSCO: assessing genome assembly and annotation completeness with single-copy orthologs. Bioinformatics, 31(19), 3210–3212.

Smith, S., Bernatchez, L., & Beheregaray, L. B. (2013). RNA-seq analysis reveals extensive transcriptional plasticity to temperature stress in a freshwater fish species. BMC Genomics, 14(1). doi:10.1186/1471-2164-14-375

Somero, G. (2010). The physiology of climate change: how potentials for acclimatization and genetic adaptation will determine ‘winners’ and ‘losers’. Journal of Experimental Biology, 213(6), 912–920.

Sorte, C. J., Jones, S. J., & Miller, L. P. (2011). Geographic variation in temperature tolerance as an indicator of potential population responses to climate change. Journal of Experimental Marine Biology and Ecology, 400(1), 209–217.

Stamatakis, A. (2014). RAxML version 8: a tool for phylogenetic analysis and post-analysis of large phylogenies. Bioinformatics, 30(9), 1312–1313.

Stillman, J. H. (2003). Acclimation capacity underlies susceptibility to climate change. Science, 301(5629), 65–65.

Summers, D. M., Bryan, B. A., Crossman, N. D., & Meyer, W. S. (2012). Species vulnerability to climate change: impacts on spatial conservation priorities and species representation. Global Change Biology, 18(7), 2335–2348. doi:10.1111/j.1365-2486.2012.02700.x

Sunday, J. M., Bates, A. E., & Dulvy, N. K. (2011). Global analysis of thermal tolerance and latitude in ectotherms. Proceedings of the Royal Society of London B: Biological Sciences, 278(1713), 1823–1830.

Sunday, J. M., Bates, A. E., & Dulvy, N. K. (2012). Thermal tolerance and the global redistribution of animals. Nature Climate Change, 2(9), 686–690.

Szklarczyk, D., Morris, J. H., Cook, H., Kuhn, M., Wyder, S., Simonovic, M., … Bork, P. (2016). The STRING database in 2017: quality-controlled protein–protein association networks, made broadly accessible. Nucleic Acids Research, 45, D362.

Tewksbury, J. J., Huey, R. B., & Deutsch, C. A. (2008). Putting the heat on tropical animals. Science, 320(5881), 1296.

Thomas, C. D., Cameron, A., Green, R. E., Bakkenes, M., Beaumont, L. J., Collingham, Y. C., … Hannah, L. (2004). Extinction risk from climate change. Nature, 427(6970), 145–148.

Thomas, C. D., & Lennon, J. J. (1999). Birds extend their ranges northwards. Nature, 399(6733), 213.

UniProt-Consortium. (2014). UniProt: a hub for protein information. Nucleic Acids Research, 43(D1), D204-D212.

Unmack, P. J., Allen, G. R., & Johnson, J. B. (2013). Phylogeny and biogeography of rainbowfishes (Melanotaeniidae) from Australia and New Guinea. Molecular Phylogenetics and Evolution, 67(1), 15–27. doi:10.1016/j.ympev.2012.12.019

Vale, C. G., & Brito, J. C. (2015). Desert-adapted species are vulnerable to climate change: Insights from the warmest region on Earth. Global Ecology and Conservation, 4, 369–379.

Vernberg, F. J. (1959). Studies on the physiological variation between tropical and temperate zone fiddler crabs of the genus Uca. II. Oxygen consumption of whole organisms. The Biological Bulletin, 117(1), 163–184.

Wellband, K. W., & Heath, D. D. (2017). Plasticity in gene transcription explains the differential performance of two invasive fish species. Evolutionary Applications, 10(6), 563–576.

Whitehead, A. (2012). Comparative genomics in ecological physiology: toward a more nuanced understanding of acclimation and adaptation. Journal of Experimental Biology, 215(6), 884–891.

Whitehead, A., & Crawford, D. L. (2006). Neutral and adaptive variation in gene expression. Proceedings of the National Academy of Sciences, 103(14), 5425–5430.

Wiens, J. A., Stralberg, D., Jongsomjit, D., Howell, C. A., & Snyder, M. A. (2009). Niches, models, and climate change: assessing the assumptions and uncertainties. Proceedings of the National Academy of Sciences, 106(2), 19729–19736.

Wiens, J. J. (2016). Climate-related local extinctions are already widespread among plant and animal species. PLoS Biology, 14(12), e2001104.

Wund, M. A. (2012). Assessing the impacts of phenotypic plasticity on evolution. Integrative and Comparative Biology, 52(1), 5–15. doi:10.1093/icb/ics050

